# Meditation induces shifts in neural oscillations, brain complexity and critical dynamics: Novel insights from MEG

**DOI:** 10.1101/2025.03.18.643795

**Authors:** Annalisa Pascarella, Philipp Thölke, David Meunier, Jordan O’Byrne, Tarek Lajnef, Antonino Raffone, Roberto Guidotti, Vittorio Pizzella, Laura Marzetti, Karim Jerbi

## Abstract

While the beneficial impacts of meditation are increasingly acknowledged, its underlying neural mechanisms remain poorly understood. We examined the electrophysiological brain signals of expert Buddhist monks during two established meditation methods known as Samatha and Vipassana, which employ focused attention and open monitoring technique. By combining source-space magnetoencephalography (MEG) with advanced signal processing and machine learning tools, we provide an unprecedented assessment of the role of brain oscillations, complexity and criticality in meditation. In addition to power spectral density (PSD), we computed long-range temporal correlations (LRTC), deviation from criticality coefficient (DCC), Lempel-Ziv complexity (LZC), 1/f slope, Higuchi fractal dimension (HFD), and spectral entropy. Our findings indicate increased levels of neural signal complexity during both meditation practices compared to the resting state, alongside widespread reductions in gamma-band LRTC and 1/f slope. Importantly, the DCC analysis revealed a separation between Samatha and Vipassana, suggesting that their distinct phenomenological properties are mediated by specific computational characteristics of their dynamic states. Furthermore, in contrast to most previous reports, we observed a decrease in oscillatory gamma power during meditation, a divergence we attribute to the correction of the power spectrum by the 1/f slope. We discuss how these results advance our comprehension of the neural processes associated with focused attention and open monitoring meditation practices.

## Introduction

Long relegated to the esoteric in the Western world, the practice and scientific study of meditation has seen a sharp rise in recent years. Meditation refers to a broad variety of practices and may be conceptualized as a family of complex emotional and attentional strategies developed for various ends, including the cultivation of well-being [42]. Over the past decades, a growing body of research has broadly supported the claim that meditation exerts beneficial effects on physical and mental health. Neuroimaging studies have begun to uncover the specific brain areas, networks and underlying attentional processes that mediate its positive effects [13, 43, 60, 68]. Expert meditators, in particular, offer a unique window into the neurodynamics associated with attention mechanisms [24, 47] and altered states of consciousness.

Emerging theories suggest that many altered states of consciousness are quantifiable through measures of criticality and complexity, which index the brain’s information processing capacity [48, 63, 72, 82]. Criticality is the dynamical state of a system that operates at the boundary of a phase transition between an ordered phase and disordered phase. The critical brain hypothesis proposes that the global neuronal dynamics of the healthy brain operate at this boundary or critical point, where emergent long-range correlations endow brain dynamics with both stability and flexibility, crucial for optimal cognitive functioning [55]. With regard to meditation and attentional processes, the ability to rapidly focus and shift attention has been linked to critical dynamics. Namely, according to a study by Irrmischer et al. [31], when the brain enters a focused task state, it appears to shift its dynamics away from criticality and into the ordered phase. The ordered phase is believed to facilitate exploitation by narrowing the range of possible responses, enhancing response stability and minimizing distractions [31]. The cited study, like many criticality studies in humans, examines critical dynamics at the level of neural oscillations [26]. While the relationship between meditation and neural oscillations has been studied extensively [38, 39, 51], research on the relation between critical brain dynamics and meditation is still at an early stage, with findings that remain unclear [17, 16, 30, 34, 76].

The concept of complexity has been variously defined, but in general it is associated with measures of entropy and fractality. A recent review by Atad et al. [3] highlighted a discrepancy in the literature on meditation and complexity, with empirical studies based on different theoretical frameworks suggesting that meditation is associated either with enhancement [34, 40, 76] or reduction of brain signal complexity [75, 80]. The clearest trends in the literature relate to expert meditators, namely that they show both (i) higher brain signal complexity [17, 34, 40, 76] and (ii) attenuated long-range temporal correlations during meditation compared to waking rest or mind wandering [17, 30, 76], as well as (iii) a reduced baseline (trait) complexity compared to novices and controls [34]. The authors also analyzed these results based on the various meditation styles, but found no differences. They suggest two possible interpretations: either complexity-related measures can characterize meditation regardless of technique, or the categorization used is too simplistic and does not accurately reflect meditators’ experiences. This categorization is centered around two main categories of meditation styles: focused attention meditation (FAM) and open monitoring meditation (OMM). FAM involves sustaining focus on a specific object (such as the breath or a specific bodily sensation), in this way enhancing concentration and attentional control [43]. An example of FAM is Samatha meditation. In OMM, on the other hand, the meditator engages in a non-specific monitoring of experiences, remaining attentive to any experience that might arise, without selecting, judging or focusing on any particular object [42, 68]. Vipassana meditation is generally assigned to this category. A number of studies have used this categorization to probe the neural correlates of meditation [1, 5, 9, 17, 29, 34, 47, 74, 75, 76, 79, 80].

Despite the growing body of research investigating neural criticality and complexity during the meditative state, the neural mechanisms underlying the different forms of meditation are still poorly understood, possibly due in part to the following factors: 1) many confounding variables, such as tradition, expertise, and control conditions, may be responsible for incoherent findings [42, 68]; 2) only a handful of studies have directly compared FAM and OMM [17, 34, 73, 74, 75, 76, 80]; and 3) much of the literature is based on fMRI [13], which lacks the temporal and spectral resolution needed to fully capture fine-grained modulations of complexity and criticality during meditation.

In the present study, we address these gaps by examining the neural dynamics of highly-trained expert Buddhist monks performing Samatha (FAM) and Vipassana (OMM) meditation, as well as in a resting state (RS), by analyzing brain signals recorded with magnetoencephalography (MEG) [2]. Combining MEG’s high temporal resolution with measures of complexity and machine learning techniques, we test the hypothesis that, compared to RS, Samatha and Vipassana are associated with distinct changes in brain signal complexity and critical dynamics. To this end, we focus on a set of complexity and criticality-related measures, namely detrended fluctuation analysis (DFA) applied to the envelope of neural oscillations, the deviation from criticality coefficient (DCC), LempelZiv complexity (LZC), spectral entropy (SpecEn), Higuchi’s fractal dimension (HFD) and the exponent of the aperiodic component of the power spectral density (PSD), also known as the 1/f slope. In tandem, we also explore changes in neural oscillations, whose role remains central to understanding brain dynamics. Previous studies have identified changes in power during meditation, particularly in the gamma range [4, 8, 41]. However, most of these studies did not distinguish between aperiodic and periodic components of the power spectrum. Given the evidence that the slope of the aperiodic component reflects the brain’s excitation-inhibition balance [20], we expect this aperiodic slope to change during meditation, potentially impacting the reliability of previous reports on spectral power alterations. Specifically, as a secondary hypothesis, we suspect that the reported increases in gamma power may be explained by a broadband shift in the aperiodic component, rather than by enhanced oscillatory activity. To address this, we use the *specparam* [15] method to separate aperiodic from periodic components of the PSD. This multifaceted approach provides an incisive and comprehensive understanding of how Samatha and Vipassana meditation modulate nonlinear brain dynamics.

## Methods and Materials

To probe the large-scale brain dynamics associated with different types of meditation, we analyzed the neuromagnetic activity in a group of expert Buddhist monks during Samatha and Vipassana meditation. More specifically, we focused on changes in spectral power, complexity and criticality-related measures.

### Participants

We recruited 12 professional meditators from the Theravada Buddhist tradition at the Santacittarama monastery, in Italy. The participants were all male, with a mean age of 38.7 years (range 25-58 years, SD 10.9 years). The monks at Santacittarama follow a Theravada Thai Forest Tradition, which includes a 3-month winter retreat meditation and a balanced practice of Samatha and Vipassana. Outside the retreat period, monks communally practice Samatha-Vipassana meditation for 2 hours daily, in addition to their own individual practice, maintaining always a balance between FAM and OMM. The participants recruited in the study had a combined meditative expertise of minimum 2375 hours (mean meditation hours 14765, range 2375–26600, SD 8018).

### Procedure

The experimental paradigm consisted in a block design of 6 min Samatha meditation and 6 min Vipassana meditation blocks, each preceded and followed by a 3 min of non-meditative resting state block (RS) [47]. Each sequence was repeated three times, for a total of 12 sessions (6 RS, 3 Samatha, 3 Vipassana). For all conditions, subjects sat in the MEG scanner with eyes closed and did not employ any discursive strategy, recitation, breath manipulation, or visualization techniques. Data were recorded using a 165-channel MEG system installed inside a magnetically shielded room at the Institute of Advanced Biomedical Technologies (ITAB), University of Chieti [12, 57]. This system includes 153 dc SQUID integrated magnetometers arranged on a helmet covering the whole head plus 12 reference channels. Electrocardiogram (ECG) and electro-oculogram (EOG) signals were also recorded for artifact rejection. All signals were band-pass filtered at 0.16–250 Hz and digitized at 1 kHz. Magnetic resonance images were acquired using a sagittal magnetization prepared rapid acquisition gradient echo T1-weighted sequence (MP-RAGE; Siemens Vision scanner 1.5 T; TR = 9.7 s, echo time TE = 4 ms, alpha = 12◦, inversion time = 1200 ms, voxel size = 1 *mm*^3^). All procedures and protocols were approved by the Ethics Committee at the University of Chieti-Pescara, and were conducted in accordance with the local legislation and institutional requirements. The participants provided written informed consent, in accordance with the Declaration of Helsinki. The same dataset has been used in [17, 47] where more details on MEG data acquisition can be found.

### Data preprocessing and MEG source reconstruction

Following standard procedures, the raw MEG data were filtered with a zero-phase band-pass using a finite impulse response filter (FIR 1, order = 3, Hamming window) between 1 Hz and 150 Hz and the filtered data were then downsampled to 600Hz. Independent component analysis (ICA) was used to remove eyeblinks, eye movements, heart-beat artifact and external artifacts from the MEG signal. These analysis steps were performed using a standard preprocessing pipeline provided by NeuroPycon (https://github.com/neuropycon), an open-source Python package that provides template pipelines for advanced and fast multi-processing of neuroimaging data [52]. We excluded two subjects from further analysis due to excessive noise in the signal and missing MRI data. Each subject’s individual MRI data was segmented using the Freesurfer software package [18] and was used to i) generate a mixed source space that includes a discretization of both the cortical surface (8196 nodes with 5 mm voxel distance) and subcortical regions (Amygdala, Caudate, Cerebellum, Hippocampus, Thalamus); ii) perform the co-registration between MRI and MEG devices; and iii) compute the forward matrix using a Boundary Element Method algorithm with a single layer model [22]. Finally, for each subject, experimental condition (RS, Samatha, Vipassana) and session, we estimated the neural time series for each voxel in the mixed source space using the weighted Minimum Norm Estimate algorithm [25]. The elemental dipoles of the cortical source-space were constrained to have an orientation normal to the surface, while the dipoles of the subcortical volumes were left with free orientation. In total, after source space morphing to a common MNI brain, we obtained 8196 time series at the cortical level, and 3 time-series per source in subcortical regions. We optimized computational efficiency by employing the source reconstruction pipeline of NeuroPycon, which runs the analyses on a multi-core processor.

### Features computation

To monitor the complexity and criticality-related dynamics of the monks’ brain activity while they were engaged in Samatha, Vipassana and eyes-closed resting-state, we extracted seven types of features from the MEG source time series: power spectral density (PSD) (both corrected and uncorrected for the 1/f spectral slope), the slope of the aperiodic component of the power spectrum, long range temporal correlations (LRTC), deviation from criticality coefficient (DCC), Lempel-Ziv complexity (LZC), Higuchi fractal dimension (HFD) and spectral entropy (SpecEn). The following sections list the spectral, complexity and criticality-related features we analyzed in this work, each computed independently for each subject, condition and session, where not otherwise specified.

#### Power Spectral Density

We computed power spectral density (PSD) at source level, in both cortical and subcortical regions, in the following frequency bands: delta (2-4 Hz), theta (5-7 Hz), alpha (8-12 Hz), beta (13-29 Hz), gamma1 (30-59 Hz) and gamma2 (60-90 Hz). We computed the PSD on the estimated neural time series using Welch’s method [77] with a window length of 4s and overlap of 2s. Importantly, to isolate the spectral components that are strictly oscillatory in nature, we removed the aperiodic 1/f-like component from the power spectrum (see the next section) by subtraction. The 1/f-corrected spectrum was then divided into the same six frequency bands defined above.

#### Aperiodic slope

We used the specparam algorithm (formerly fooof) [15] to estimate the slope of the aperiodic component of the estimated neural time series, which corresponds to the exponent of the 1/f-like distribution of the power spectrum. The specparam algorithm decomposes the log power spectra into a summation of narrowband gaussians (periodic oscillations) and a linear trend (aperiodic component) within a broad frequency range. We used a 1-90 Hz frequency range and limited the number of oscillatory peaks to 6. In the following analysis, we express the slope as the exponent *α* in 1*/f^α^*, such that an increase of the slope corresponds to a steeper decline in power. EEG modelling work has suggested that such steepening may be explained by an increase in global cortical inhibition [20], but see [6].

#### Long-range temporal correlations

We investigated the temporal dynamics of neural oscillations by probing for the presence of long-range temporal correlations (LRTC) in the amplitude envelope of the oscillations in different frequency bands using Detrended Fluctuation Analysis (DFA) [26, 54]. DFA is a well-established method that measures the fluctuations of a signal as a function of a sliding time-window at multiple scales. The original signal is integrated to create a cumulative sum series and split into windows of different sizes that are equidistant on a logarithmic scale. In each window, the linear trend is removed and the fluctuation function is defined as the average root-mean-square for each window size. The slope of the line fitting the fluctuation function in the log-log plot is the LRTC exponent and gives an indication of the scaling behavior of the time series. It is often referred to as the self-similarity parameter and can take on a value between 0 and 1, where a value between 0 and 0.5 represents negative temporal correlation (i.e. anti-persistence of the signal in time), 0.5 represents a random (time series is uncorrelated) signal, and a value between 0.5 and 1 represents a positive temporal correlation. In a persistent signal, large signal values are likely to be followed by more large signal values and small values are likely to be followed by more small values. Conversely, in an anti-persistent signal (negative correlation) large values are likely to be followed by small ones and small values are likely to be followed by large ones. Thus, if the LRTC exponent decreases from a value close to 1 down to a value closer to 0.5, then the temporal correlations are considered to be less persistent in time (i.e. they decay faster in time). We performed DFA on the amplitude envelope of the neural oscillations in alpha, beta, gamma1 and gamma2 bands. To this end, we filtered the estimated neural time series using a finite impulse response filter (FIR1, order = 3), and applied the Hilbert transform [19, 38] to extract the amplitude envelope. DFA values were computed for each of the three conditions (RS, Samatha, Vipassana), in the time range of 2–50 s over consecutive windows (no overlap). To render the calculation of the DFA more robust, we concatenated the different sessions of each condition. As highlighted in [26], DFA estimates for window sizes *>*10% of the signal length are noisier due to a low number of windows available for averaging (cf. [26, 30]).

#### Deviation from criticality coefficient

Neural activity can at many scales be observed to spread as cascades through space and time, known as neuronal avalanches [55]. At the critical point, neuronal avalanches exhibit scale invariance; that is, the probability distribution of various avalanche properties, such as their size and duration, follow a power law. While the detection of such power-law behavior provides an indicator of criticality, on its own it is not a very specific indicator, as other simple processes also yield such power-law scaling [71]. However, criticality theory predicts that the exponents of these power laws should follow certain interrelations, known as scaling relations [53], that are not expected to be obeyed away from criticality [71]. The degree of adherence to these relations is thus a fairly reliable indicator of the distance to criticality. One of these relations, known as the crackling noise relation [65], connects the critical exponents for the distributions of avalanche size S (the total number of events, i.e. the sum of events across all voxels), avalanche duration T, and average S for every T:

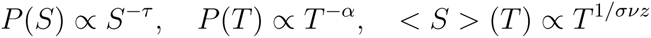

which should be related as:

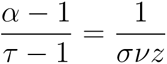

We note that this relation holds generally for any class of criticality with an absorbing state [53]. From this relation, the deviation from criticality coefficient (DCC) ([44]) is defined as:

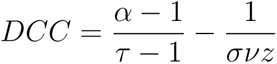

To detect neuronal avalanches, source-reconstructed time series were z-scored voxelwise and binarized according to a threshold determined using the following data-driven method [67, 72]. The z-scored time series were first averaged across all subjects within the same condition and plotted in a probability distribution ranging from −10 to 10 SD. A Gaussian was fitted to this distribution and the binarization threshold was taken as the point where the Gaussian diverges from the data distribution, which represents the SD at which neural events are no longer expected to be the result of merely stochastic fluctuation. Following this method, the binarization threshold was set to ±3 SD and negative and positive excursions beyond the threshold were identified as events and assigned a value of one at their maximum/minimum value. An avalanche was defined as a continuous sequence of events, on any voxel, separated in time by at most the time bin *t* = 4 ms [67]. Avalanches were detected in this way for each subject and condition. The size, duration, and average size by duration distributions were plotted, and the critical exponents fitted from the distributions. From these exponents, we computed the DCC for each ROI of the Yeo 17-network atlas [78] using the *edgeofpy* Python package (https://github.com/jnobyrne/edgeofpy).

#### Lempel-Ziv complexity

Lempel-Ziv complexity (LZC) provides a measure of entropy by counting the number of distinct patterns of activity in the data [38]. It can be thought of as being proportional to the size of a computer file containing the data, after applying a compression algorithm [64]. LZC is a complexity measure designed for binary sequence and text and is computed on a binary version of the signal. The time series were binarised by using the median value: the values above the median are assigned ones, and below the median are assigned zeros. LZC is defined as the number of unique subsequences in the whole binary sequence. A less complex signal consists of repetitions of a few different sub-strings while more complex signals are made up of non-repeating segments. As the original version of LZC is strongly influenced by signal length, we used a normalized version which scales LZC by log*_b_*(*n*)*/n*, where n is signal length and b the number of unique characters in the signal [81].

#### Higuchi Fractal dimension

We used Higuchi’s algorithm [28] to compute the fractal dimension of the estimated neural time series. From the original time series X, k new different time series *X^k^, m* = 1*, .., k* are constructed and their average length *L*(*k*) is computed (m indicates the initial time sample and k denotes the time interval). This step is repeated for *k* = 1*, …, k_max_*. The average length *L*(*k*) is proportional to *k^−D^*, then plotting *log*(*L*(*k*)) against *log*(1*/k*) the slope of the line fitting the points in the log-log plot represents the Higuchi fractal dimension (HFD). Numerical values of HFD have the lower and upper limits of 1 and 2, respectively. Higher values of the slope indicate a more complex curve, whereas values close to 1 indicate a simple curve with low complexity. HFD can be imaged as a measure of the ‘degree of filling out’ of the plane by the curve and hence, its complexity [37]. The value of *k_max_*, the maximum number of subseries composed from the original time series, was determined by examining the data and plotting the fractal dimension over a range of k. For k greater than *k_max_* the fractal dimension reaches a saturation point [35, 45, 58, 59]. Based on this procedure, we found the optimal *k_max_* value to be 10. We also tested the robustness of the method with respect to the length of the time series and different values of k.

#### Spectral Entropy

To quantify the complexity of the power spectrum, we computed the spectral entropy (SpecEn) [56] by computing the Shannon entropy of the PSD of the neural time series. Shannon entropy [66] measures the uncertainty associated with a random variable and is calculated by taking the average value of the logarithm of the probability of each possible event. The power spectrum we used to compute SpecEn was estimated as described above, except that it was neither corrected for the 1/f-like component nor split into frequency bands.

### Statistical analysis and machine learning

We examined the task-based modulations of all computed features (PSD, slope, LRTC, LZC, HFD, SpecEn) using (a) group-level differences assessed using cluster-based permutation tests [46] and (b) a multi-feature supervised learning approach, where a Random Forest (RF) classifier was trained to discriminate between the different brain states in a k-fold cross-validation scheme (see below). We assessed the statistical differences in the DCC across the three conditions using the non-parametric Wilcoxon signed-rank test, applied to each ROI within the Yeo 17 network.

#### Cluster-based permutation test

For each feature, statistical analysis was conducted to look for group-level differences in 1) in each meditative state (Samatha, Vipassana) as compared to RS condition and 2) between the two meditative states. Since for each subject we used an individual source space, before performing the statistical hypothesis test we morphed and projected individual source space onto the average brain *fsaverage* provided by Freesurfer.

Differences between the different conditions (RS, Samatha, Vipassana) for the several features were assessed using the cluster-based permutation tests corrected for multiple comparisons as implemented in MNE-python [23]. The procedure uses a cluster analysis with permutation test for calculating corrected p-values. First, clusters are identified based on the spatial adjacency of voxels where the observed effect (difference between conditions assessed by a paired t-test) is statistically significant (*p_val_ <* 0.05). Then, each cluster is associated with a cluster level statistic (the sum of voxel t-values within the cluster) that is compared to the null distribution of 1024 permutations using shuffled labels. Differences between conditions are considered significant when the observed cluster-level statistic exceeds the maximum cluster level statistics from the null distribution using a threshold of p-value = .05 corrected for multiple comparisons [46].

#### Pearson correlation analyses

We evaluated the relationship between hours of meditation practice and the features where we found a state effect induced by the meditative condition. Specifically, we calculated the relative difference (*M* − *RS*)*/RS* of the features, where M is either Samatha or Vipassana. This was computed within voxels of the clusters determined to be significant in the previous analysis. Individual values in these clusters, for each subject, were averaged across each cluster and then used to calculate their Pearson correlation with the expertise of each monk. To this end, we used the log of the total hours of meditation, as in previous reports [5]. Additionally, for each meditation condition, we computed the Pearson correlation between feature pairs, considering for each pair the average values across common voxels, as defined by the significant clusters.

#### Multi-feature classification

We additionally implemented a multi-feature classifier to characterize the contribution of the different features in discriminating between meditative states and resting state condition. To identify which brain regions contribute more to the separation we reduce each feature to a low dimensional space by computing the mean values over the regions of interest (ROI) provided by the 450-label aparc-sub parcellation [36] and the 10 sub-cortical regions. Then, we took together all features from all ROIs to construct a single model and we were able to access the contributions of individual features and ROIs by examining feature importance. To achieve this we used a random forest [7], which estimates feature importance by the relative rank of each feature across the decision trees that make up the forest. To measure variance in feature importance, the training process was repeated 120 times for each comparison (Samatha vs RS, Vipassana vs RS, Samatha vs Vipassana), applying grid search with grouped 7-fold cross-validation inside a nested cross-validation, leaving out samples from three subjects in each iteration. The overall model score was determined using samples from left-out testing subjects and the Area Under the Curve (AUC) was used to evaluate the performance of the classifier [69].

## Results

The following section describes the results obtained by contrasting the three different conditions (RS, Samatha and Vipassana). We start by reporting spectral data, which provide insight on changes in periodic brain activity, then we describe the results of non-parametric statistical tests for each of the extracted features (narrowband power, 1/f slope, LRTC, LZC, HFD, SpecEn and DCC) and the output of the multi-feature classifier. Except for DCC (Figures 5) the direct comparison between Samatha and Vipassana did not yield significant differences. The summary of these findings for the non-significant features (narrowband power, 1/f slope, LRTC, LZC, HFD, SpecEn) is provided in the supplementary material (Fig. S1).

### Meditation induces changes in MEG oscillations

Figures 1A and 2A show the results of the source-level spectral power analysis by frequency band, for Samatha vs. RS and Vipassana vs. RS, respectively. The left column shows the spatial distribution of t-values obtained by two-tailed paired t-tests, while the right column displays only the t-values that were significant (*p <* 0.05) based on cluster-based permutation tests. Samatha meditation led to a statistically significant reduction of power (*p <* 0.05) in beta, gamma1 and gamma2 bands, mainly in occipital and parietal regions.

**Figure 1:**
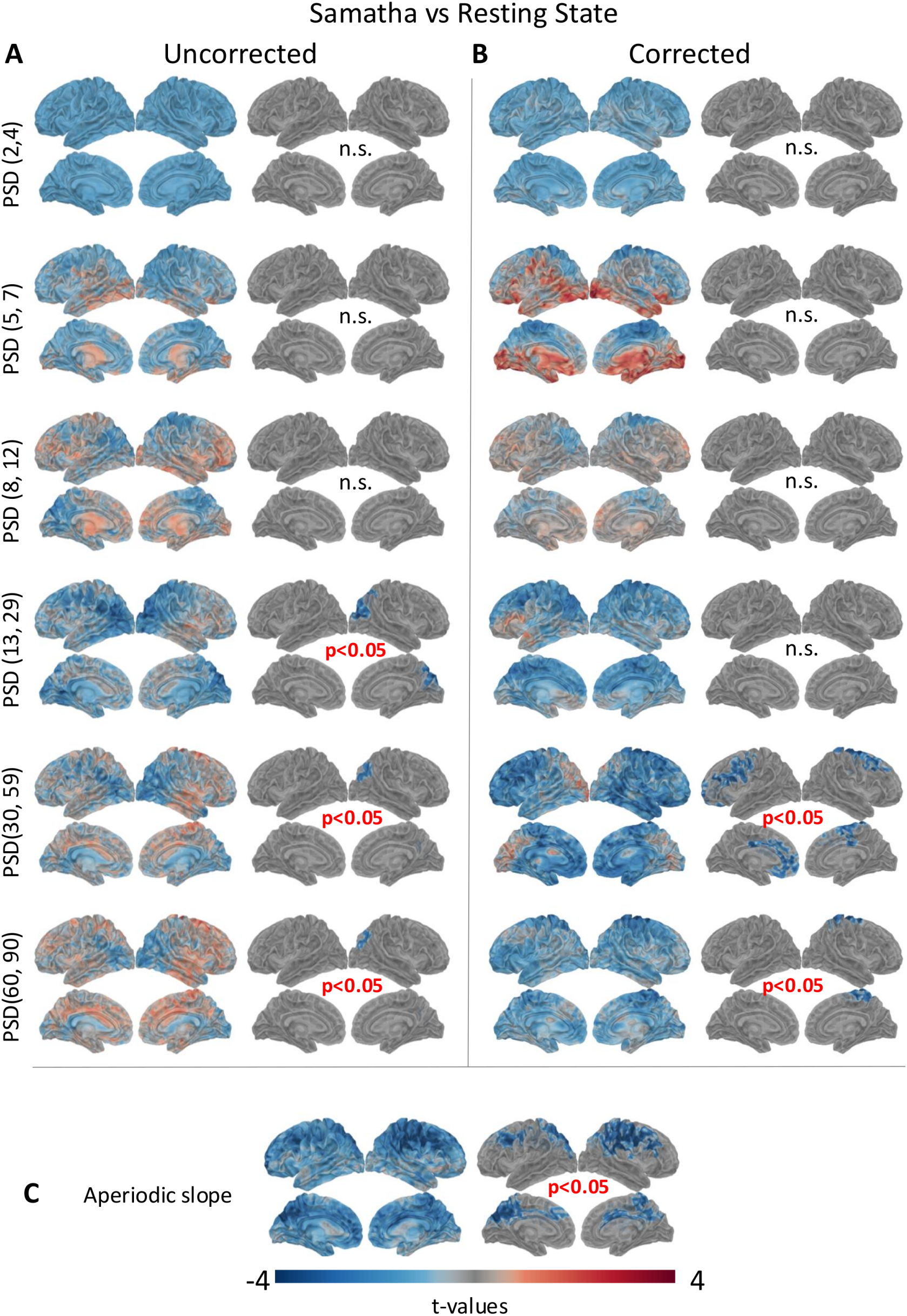
Results of the source-level spectral power analysis in the different frequency bands for Samatha as compared to RS condition. A) The left column shows the results of two-tailed paired t-tests, and the right column shows the results of cluster-based permutation tests (*p <* 0.05). B) T-maps and results of cluster-based permutation tests for spectra corrected for the 1/f slope. C) T-maps and results of cluster-based permutation tests for the aperiodic component (1/f slope) of the power spectra.

**Figure 2:**
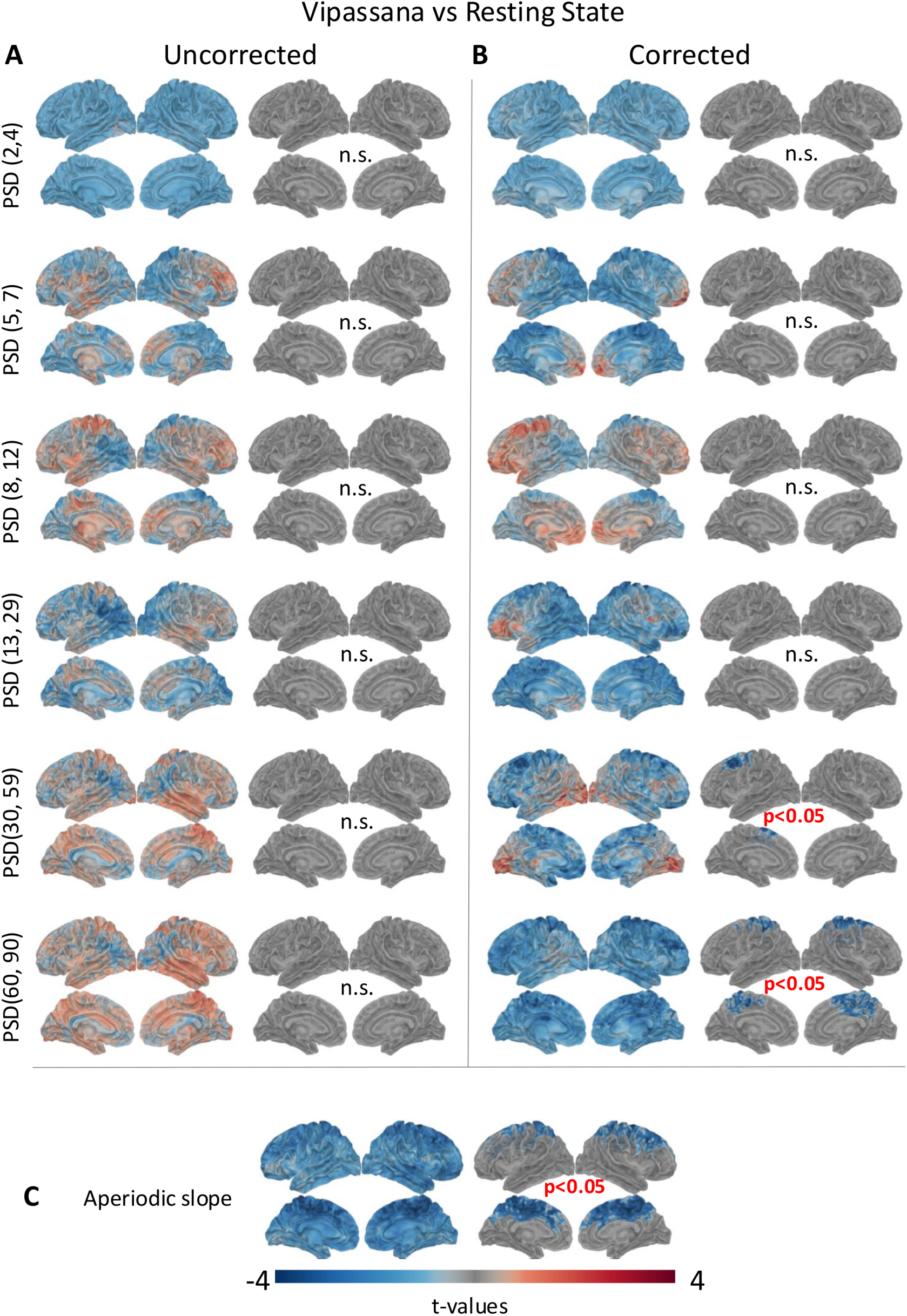
Results of the source-level spectral power analysis in the different frequency bands for Vipassana as compared to RS condition. A) The left column shows the results of two-tailed paired t-test, and the right column shows the results of cluster-based permutation tests (*p <* 0.05). B) T-maps and results of cluster-based permutation tests for spectra corrected for the 1/f slope. C) T-maps and results of cluster-based permutation test for the aperiodic component (1/f slope) of the power spectrum.

Interestingly, the spectral results are quite different when we correct for the aperiodic slope (Figures 1B, 2B). Because the power spectrum is the sum of peaks of oscillatory (periodic) activity and a background of non-oscillatory (aperiodic) frequency content [15], not performing the correction may lead to confounding results that conflate oscillatory and non-oscillatory signals [61]. With the aperiodic component removed in our data, we note the direction of the effect is consistent for all frequency bands, with the exception of the beta band for Samatha and the gamma bands for Vipassana. Discarding the aperiodic component reveals that both meditation practices induced a statistically significant reduction in gamma power (*p <* 0.05), especially in the precentral and superior frontal regions (Figure 1B, 2B right column).

Figures 1C and 2C show the t-maps and the results of cluster-based permutation tests for the aperiodic slope for Samatha vs. RS and Vipassana vs. RS, respectively. We found a statistically significant reduction of the slope (*p <* 0.05) in both meditative conditions, mainly in the frontal, parietal, occipital and temporal regions. This finding supports our choice to perform the correction. Interestingly, we also observed a remarkably consistent relationship when comparing slopes between the different conditions across ROIs, whereby the slope was always highest during RS and lowest during Vipassana, with Samatha occurring in between (Figure 3). Such changes in the scaling behavior of the power spectra suggest meditation-induced changes in self-similarity of the electrophysiological signal, which may be related to an alteration in neural criticality. In addition, previous work supports the notion that a reduction in the 1/f exponent (i.e. flatter slope) may reflect an increase in the E-I ratio.

**Figure 3:**
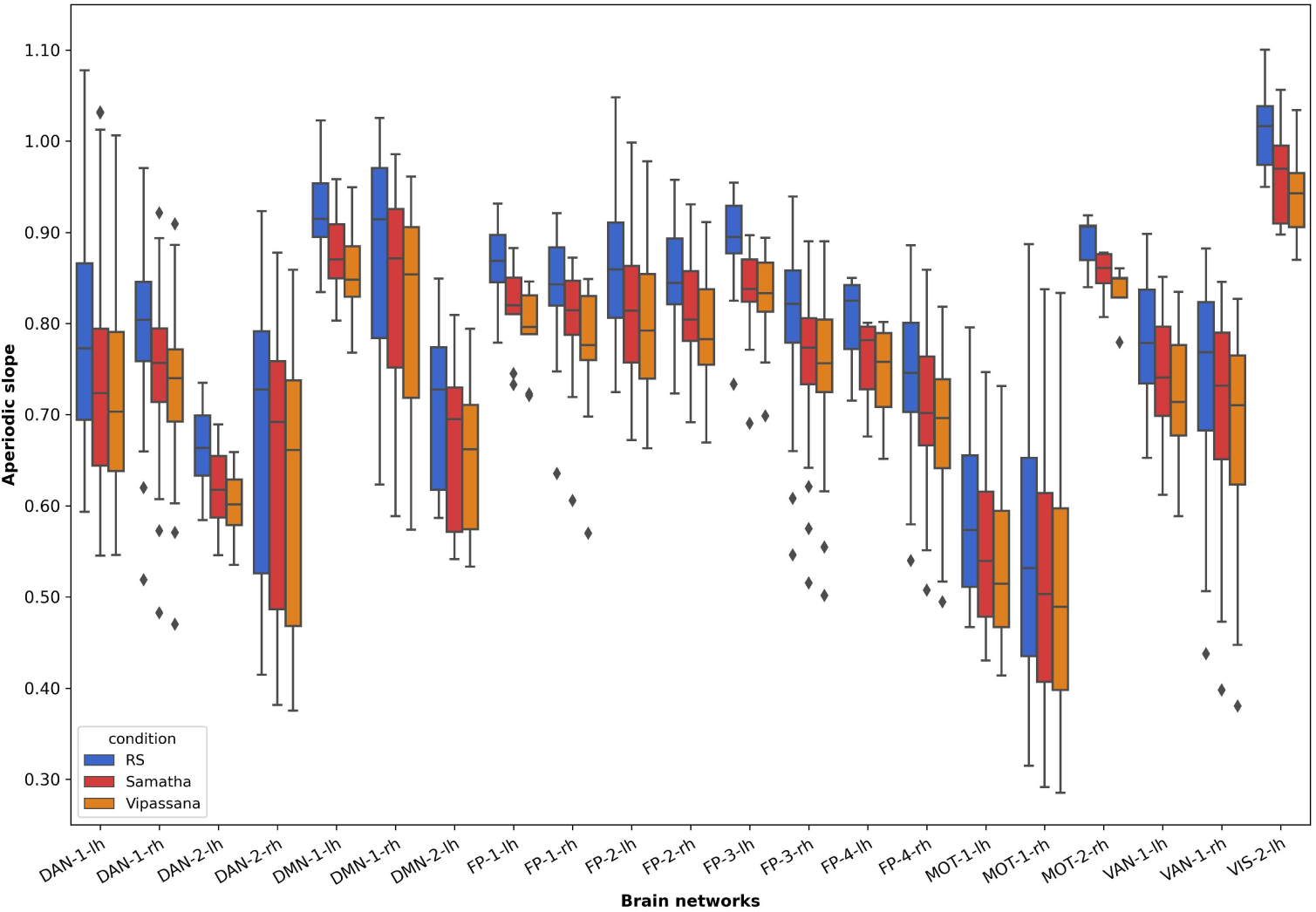
Box plot of the aperiodic slope for the different conditions: RS, Samatha and Vipassana. Each ROI belongs to the Yeo 17 networks atlas [78] and the corresponding box plots represent the values in the voxels where there is a statistical difference for both meditations with respect to the RS, averaged across subjects. The diamonds represent the outliers of data distribution.

### Meditation alters complexity and criticality-related measures

We investigated the effects of meditation on the temporal persistence of neural oscillations by computing the LRTC exponent in different frequency bands (alpha, beta, gamma1 and gamma2). Across conditions, we consistently found values for the LRTC exponent *>* 0.5, indicating a persistent, positively correlated signal. The t-maps obtained by the two-tailed paired t-tests and the results of the cluster-based permutation tests (*p <* 0.05) are shown in the top half of Figure 4 for Samatha (left) and Vipassana (right). The t-value maps reflect the magnitude of the effect and the direction of the group differences for the LRTC values, with negative (blue) t-values indicating the brain areas where meditation induced a reduction of the LRTC exponent. Marked decreases in LRTC were found across the cortex for gamma1 and gamma2 bands, in both meditation conditions. The reduction in LRTC values represents the lower intrinsic fluctuations in neural activity that indicates a shift from more complex brain dynamics to a state of reduced information propagation and reduced temporal complexity of oscillations [30]. This result is in line with the decreasing slope of the aperiodic activity: both metrics show consistent reductions and may be indicative of a shift in critical behavior.

**Figure 4:**
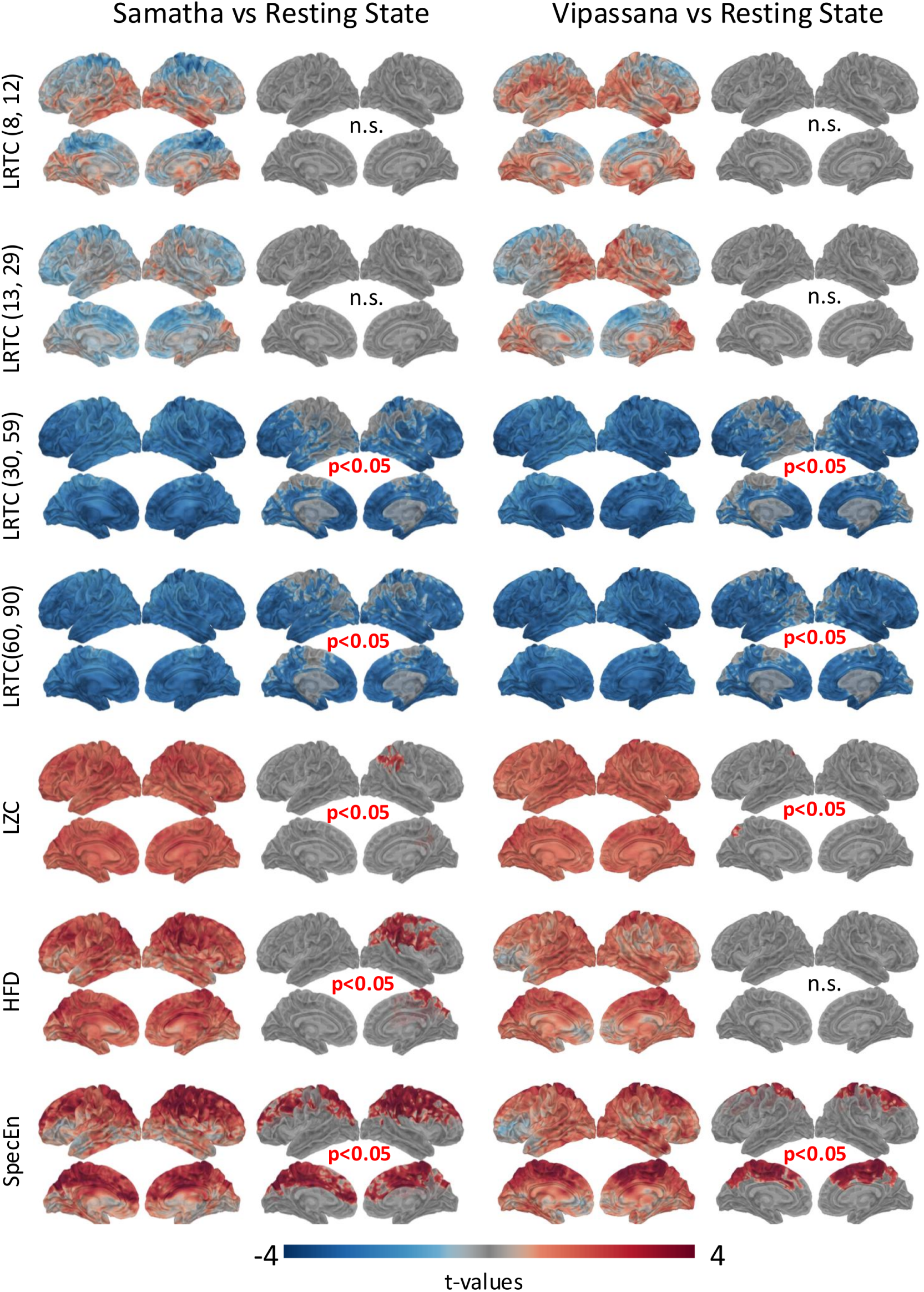
T-maps and results of cluster-based permutation tests for the different features (LRTC, LZC, HFD and SpecEn) when the two meditation conditions (Samatha on left, Vipassana on right) are compared to the RS condition.

In addition, we found a consistent trend of decreasing DCC (i.e. distance from the critical point) from RS to Vipassana and from Samatha to Vipassana (Figure 5). This decrease was statistically significant (p*<*0.05, uncorrected) in specific ROIs within the Ventral Attention, Dorsal Attention, Motor, and Default Mode Networks (Table 1).

**Figure 5:**
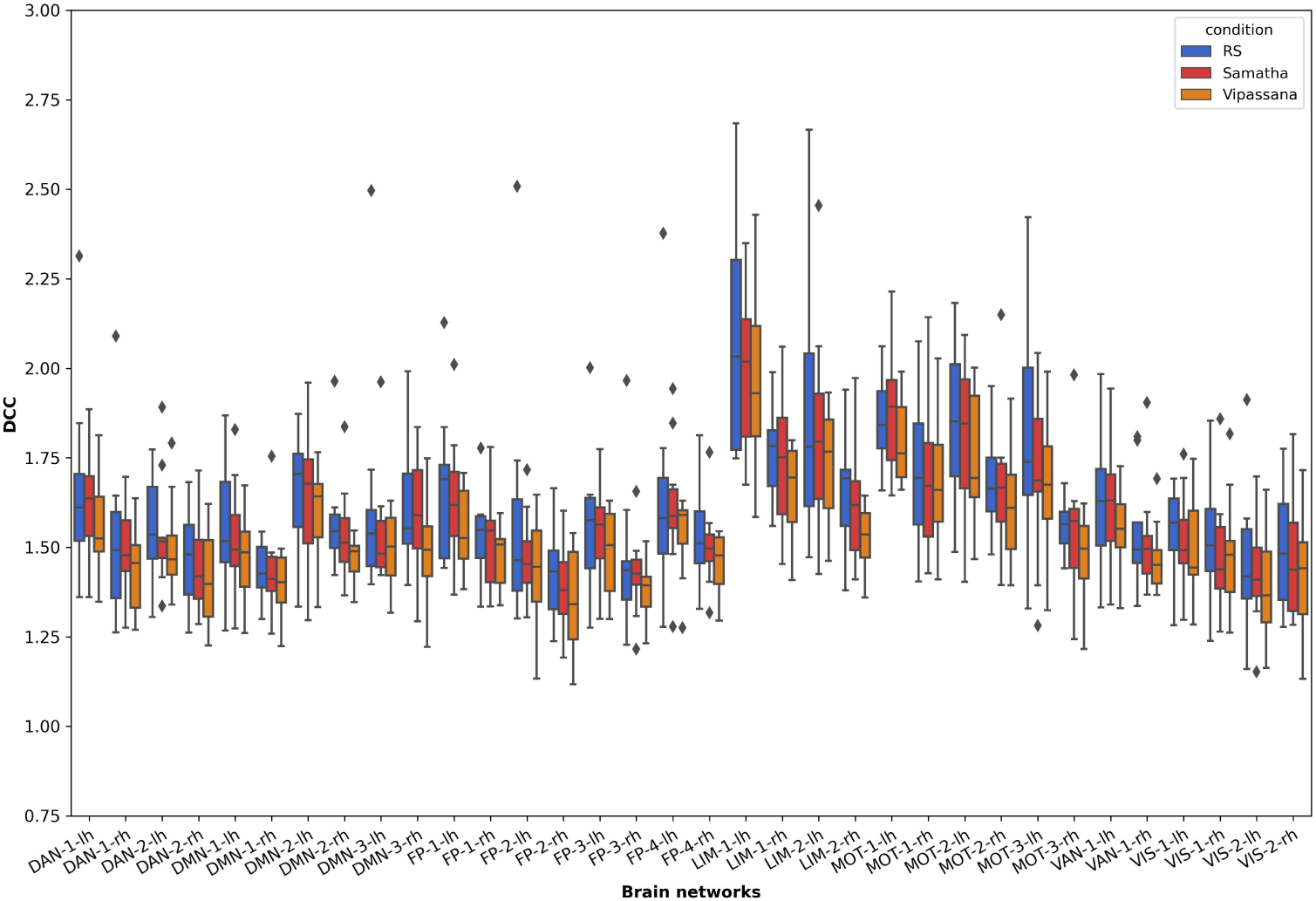
Box plot of the Deviation from criticality coefficient (DCC) for the different conditions computed for each ROI of the Yeo 17 networks atlas [78].

**Table 1:** P–values from the non-parametric Wilcoxon signed-rank test for each ROI in the Yeo 17 network, assessing pairwise comparisons of DCC across the three conditions.

| ROI | Rs vs Vipassana | Samatha vs Vipassana |
| --- | --- | --- |
| LIM-1-rh | 0.049 | 0.028 |
| LIM-2-rh | 0.006 | 0.02 |
| FP-3-rh | 0.028 | n.s. |
| FP-4-rh | 0.038 | n.s. |
| MOT-2-lh | 0.049 | n.s. |
| MOT-2-rh | n.s. | 0.0098 |
| MOT-3-rh | 0.014 | 0.009 |
| DAN-1-lh | n.s. | 0.048 |
| VAN-1-lh | n.s. | 0.014 |
| VAN-1-rh | n.s. | 0.014 |
| DMN-3-rh | n.s. | 0.049 |
| DMN-2-rh | 0.038 | n.s. |

The LZC and HFD t-maps show an overall increase in both metrics (Figure 4) for the two meditation conditions as compared to RS. In particular, we note a significant increase (*p <* 0.05) of LZC in Samatha mainly in parietal and temporal regions of the right hemisphere, and in Vipassana in precuneus and parietal regions of the left hemisphere. For HFD, the results of cluster-based permutation tests reveal a significant increase (*p <* 0.05) only for Samatha in the right hemisphere, mainly in the precentral, postcentral, superior temporal and frontal regions.

Finally, the SpecEn feature (Figure 4) quantifying the complexity of the power spectra significantly increased (*p <* 0.05) in both meditation conditions, mainly in the cingulate, central and frontal regions.

### Correlation analysis

Figure 6 shows both the significant correlations between the computed features and the hours of meditation, as well as the box plots for the values of the individual features (averaged within the significant clusters) for both main comparisons: Samatha vs. RS and

**Figure 6:**
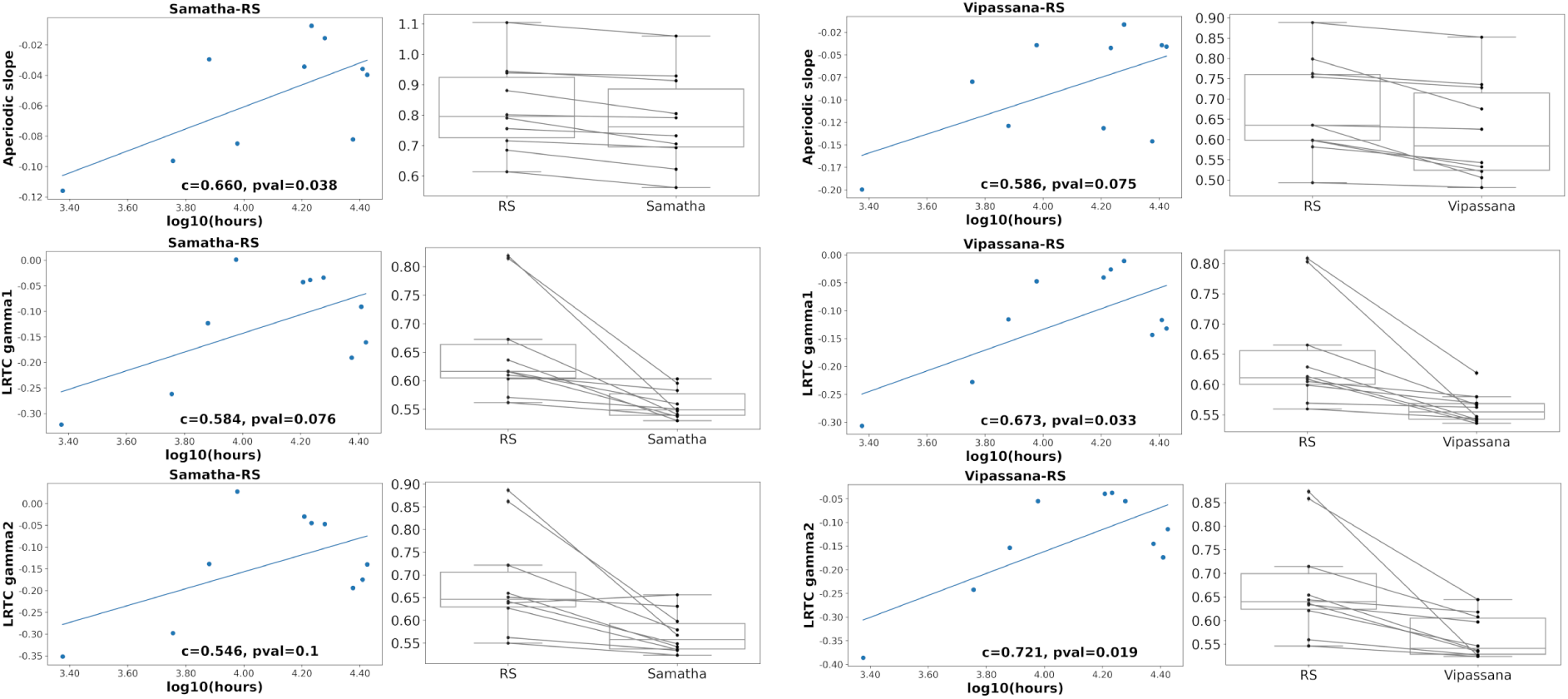
Correlations with hours of meditation and box plots, by feature. The first and third columns show the Pearson correlation between hours of meditation and the features in which we found a state effect of Samatha and Vipassana, respectively. The second and fourth columns illustrate the box plot of individual features values (averaged within the significant clusters) for Samatha and Vipassana, respectively.

Vipassana vs. RS. The number of hours of Samatha meditation was significantly positively correlated with the aperiodic slope (p=0.038) while hours of Vipassana meditation showed a significant positive correlation with the DFA exponent for gamma1 (p=0.033) and gamma2 (p=0.019). The boxplots (Figure S2) show, for both meditation conditions, a local state effect characterized by a decrease in spectral features (PSD gamma1, PSD gamma2) and criticality-related measures (aperiodic slope, DFA gamma1, DFA gamma2), as well as an increase in complexity measures (LZC, HFD and SpecEn).

Finally, Figure S3 shows, for each meditation condition, the correlations between feature pairs across common voxels in significant clusters and Figure S4 shows the correlation map between DCC and all other features computed across all conditions in each ROI of the Yeo 17 network.

### Multi-feature Machine learning

We built a Random Forest classifier that we trained on all extracted features (14 measures x 460 ROIs = 6440 total) giving us an insight into individual feature importance scores by ranking and comparing the contributions of the different features in discriminating between meditative states and RS condition. For the PSD features, we considered only those corrected for the aperiodic component. We found that the multi-feature RF classifier achieved a higher decoding accuracy for Samatha vs. RS (AUC=92.18 ± 0.12) and Vipassana vs. RS (AUC=91.67±0.1) than for Samatha vs. Vipassana (AUC=49.5±0.16). To assess the significance of the obtained AUCs, we performed a permutation test for each comparison by shuffling the feature labels 100 times and computing the 95th percentile of the null distribution. We found that observed AUC for Samatha and Vipassana were statistically significant.

Figure 7 displays the ranked feature importance, colored according to the feature category. The features of highest importance, appearing towards the left-hand side of the plots, were the ones most relevant in discriminating between the different brain states. For Samatha and Vipassana vs. RS, the most important features were the DFA exponents in gamma1 and gamma2 bands (in blue). The same was true for Samatha vs. Vipassana, with the addition of alpha-band DFA exponents. The cortical maps (inset) represent the spatial distribution of the 20 most important measure-ROI pairs (all of which happen to be in DFA alpha, DFA gamma1 or DFA gamma2). Here, we show the average feature importance over different runs: higher values indicate higher importance of a measure at a specific ROI. The ROIs that contributed the most in classifying the meditation conditions vs. RS were located mainly in the frontal, anterior cingulate, lingual and insular regions for Samatha, and in the superior frontal, precentral and parietal regions for Vipassana. Table 2 provides a list of these 20 important measure-ROI pairs as determined by the RF classifier. Overall, these results highlight DFA, a criticality-related measure, as highly discriminative between meditation and RS, as well as between different meditation styles.

**Figure 7:**
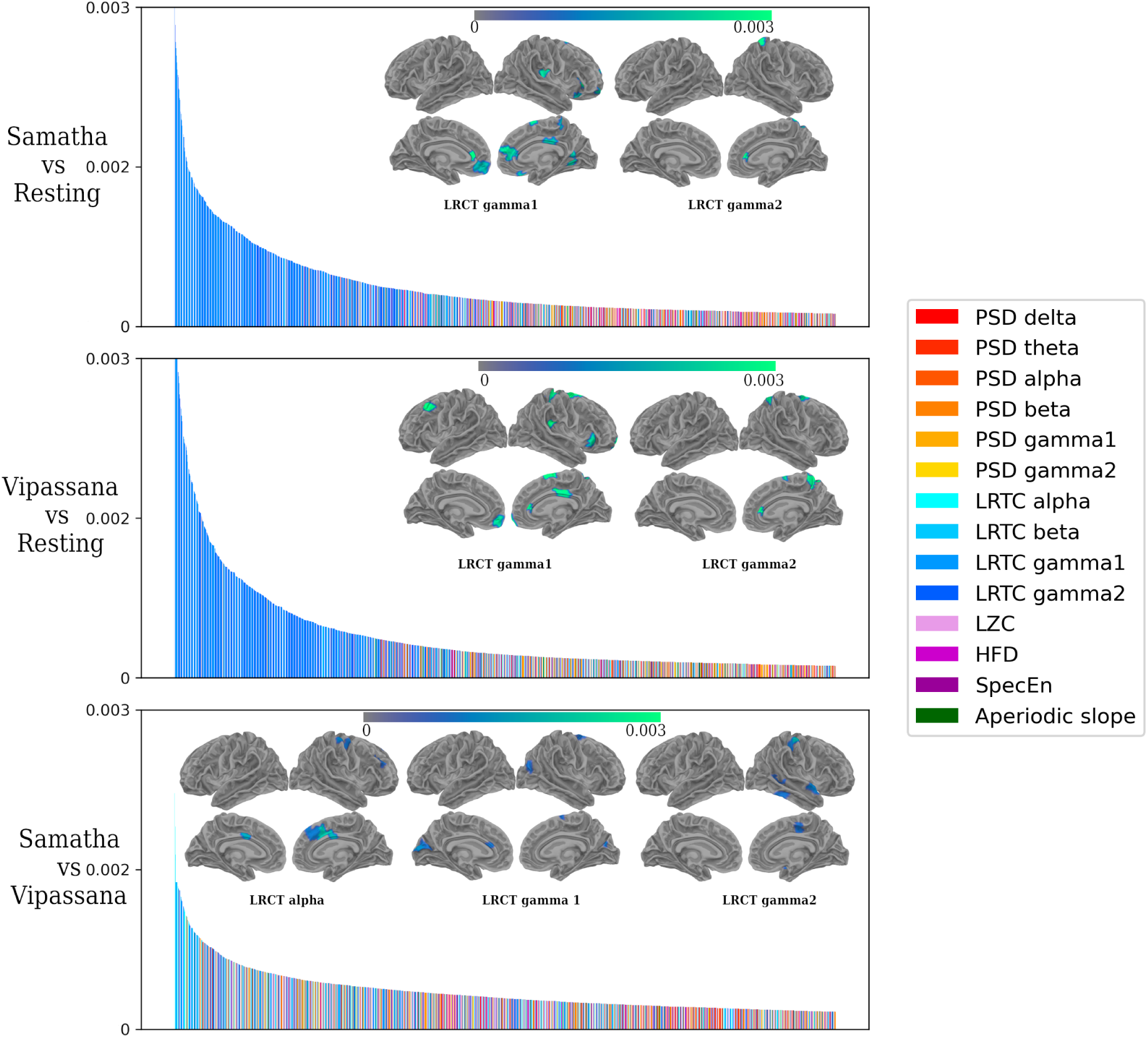
Visualization of ranked feature importance for the random forest classifiers trained on a feature space which combined all measures across ROIs (14 measures x 460 ROIs = 6440 total), color-coded according to the corresponding measure. Warm colors (red to yellow) show spectral power in the different frequency bands, while the other colors (blue, violet and green) correspond to complexity and criticality-related measures. The cortical maps depict the spatial distribution of 20 most important features. Here we show the average feature importance over different runs: higher values indicate higher importance of a feature at a specific ROI.

**Table 2:** List of the 20 most important measure-ROI pairs as determined by the Random Forest classifier.

| Samatha vs Resting State |  |  |  | Vipassana vs Resting State |  |  |  |
| --- | --- | --- | --- | --- | --- | --- | --- |
|  | feature | ROI | importance |  | feature | ROI | importance |
| 1 | LRTC gamma1 | parstriangularis_1-rh | 0.003208 | 1 | LRTC gamma1 | superiorfrontal_15-rh | 0.004914 |
| 2 | LRTC gamma1 | rostralanteriorcingulate_2-lh | 0.003206 | 2 | LRTC gamma2 | superiorparietal_4-rh | 0.004251 |
| 3 | LRTC gamma2 | rostralanteriorcingulate_2-rh | 0.002893 | 3 | LRTC gamma1 | precentral_14-rh | 0.003991 |
| 4 | LRTC gamma2 | superiorparietal_3-rh | 0.002833 | 4 | LRTC gamma1 | postcentral_11-rh | 0.003598 |
| 5 | LRTC gamma1 | superiorfrontal_15-rh | 0.002674 | 5 | LRTC gamma1 | frontalpole_1-rh | 0.003543 |
| 6 | LRTC gamma1 | superiorfrontal_3-rh | 0.002616 | 6 | LRTC gamma1 | caudalmiddlefrontal_2-lh | 0.003346 |
| 7 | LRTC gamma1 | Hippocampus-lh | 0.002544 | 7 | LRTC gamma1 | superiorfrontal_17-rh | 0.003220 |
| 8 | LRTC gamma1 | insula_1-rh | 0.002481 | 8 | LRTC gamma1 | posteriorcingulate_2-rh | 0.003108 |
| 9 | LRTC gamma1 | superiorfrontal_2-rh | 0.002414 | 9 | LRTC gamma2 | rostralanteriorcingulate_2-rh | 0.003030 |
| 10 | LRTC gamma1 | lingual_6-rh | 0.002383 | 10 | LRTC gamma1 | supramarginal_9-rh | 0.003006 |
| 11 | LRTC gamma1 | rostralmiddlefrontal_13-rh | 0.002378 | 11 | LRTC gamma1 | rostralanteriorcingulate_2-rh | 0.002890 |
| 12 | LRTC gamma2 | superiorparietal_4-rh | 0.002363 | 12 | LRTC gamma1 | postcentral_10-rh | 0.002874 |
| 13 | LRTC gamma1 | medialorbitofrontal_4-rh | 0.002344 | 13 | LRTC gamma1 | superiorparietal_4-rh | 0.002872 |
| 14 | LRTC gamma1 | posteriorcingulate_3-rh | 0.002236 | 14 | LRTC gamma1 | postcentral_9-rh | 0.002834 |
| 15 | LRTC gamma1 | precuneus_2-rh | 0.002227 | 15 | LRTC gamma2 | precuneus_9-rh | 0.002768 |
| 16 | LRTC gamma1 | medialorbitofrontal_1-lh | 0.002146 | 16 | LRTC gamma2 | superiorparietal_6-rh | 0.002664 |
| 17 | LRTC gamma1 | rostralanteriorcingulate_2-rh | 0.002128 | 17 | LRTC gamma2 | superiorfrontal_17-rh | 0.002624 |
| 18 | LRTC gamma1 | lateralorbitofrontal_3-rh | 0.002105 | 18 | LRTC gamma1 | posteriorcingulate_3-rh | 0.002623 |
| 19 | LRTC gamma1 | paracentral_1-rh | 0.002047 | 19 | LRTC gamma1 | medialorbitofrontal_1-lh | 0.002549 |
| 20 | LRTC gamma1 | medialorbitofrontal_2-lh | 0.002021 | 20 | LRTC gamma1 | lateralorbitofrontal_2-rh | 0.002545 |

## Discussion

While meditation continues to garner widespread interest and becomes more integrated into therapeutic practices, the electrophysiological changes it induces in the brain remain poorly understood. Yet, a better understanding of the neural dynamics underlying distinct meditation techniques is needed in order to tailor interventions for specific psychological conditions, optimize therapeutic outcomes, and refine methodologies for training and practice. Here, we analyzed rare neuromagnetic data recorded in expert meditators while they performed two types of meditation known as Samatha and Vipassana. Our analysis extends previous work not only by leveraging source-space MEG analyses, but also by using state of the art methods from dynamical systems theory, complexity science, spectral analyses and machine learning. First of all, we examined the effect of separating periodic and aperiodic components of the power spectrum on meditations-induced spectral power changes. This allowed us to probe whether previously observed increases in gamma power during meditation could in fact be attributed to a broadband upward shift in the aperiodic component in the higher frequencies, rather than actual increases in gamma oscillation amplitude. Secondly, we focused on characterizing changes in non-linear brain dynamics induced by the different meditation techniques through the lens of signal complexity and criticality-related metrics. In the subsequent sections, we delineate our principal findings, contextualizing them within the framework of existing literature. Additionally, we acknowledge the constraints of our study and outline prospective directions for further research.

### Meditation induces changes in spectral power and 1/f slope

Our findings on power spectral density reveal a reduction in the beta-band power for Samatha, in line with other studies using experienced meditators [1, 14, 29, 62], which may reflect increased cortical activation of sensory-related attentional networks and an increased capacity of the expert meditators to focus their attention on their breath.

Interestingly, when we used 1/f-corrected power spectra (i.e. excluding the aperiodic component), the results revealed that, compared to rest, both meditation conditions exhibited a decrease in oscillatory gamma power. An intriguing finding, given previous reports of increases in gamma-range activity during meditation (see discussion below).

High-frequency gamma oscillations have been shown to be involved in attentional mechanisms subserving visual processing and working memory [32, 33]. Therefore, the decrease in oscillatory gamma power we found here could reflect a reduction in the mental activity that arises from processing external stimuli and engaging diverse cognitive processes. This aligns with the meditative goal of achieving a state of mental calm and inward focus. The brain areas in which we observed a statistically significant reduction are mainly the frontal, parietal, and cingulate regions, which are key regions in several brain networks, including the Frontoparietal, Dorsal Attention, Motor and Default Mode networks [78]. Interestingly, the observed reduction in gamma power across these brain regions is putatively consistent with previous reports of decreases in fMRI activation in the default-mode network [21], and with the Topographic Reorganization Model of Meditation (TRoM) proposed by Cooper and colleagues [13]. The proposed framework, based primarily on fMRI findings, explains how advanced meditation reorganizes brain topography promoting a shift away from cognitive engagement toward a state of integrated awareness, associated

The observed decrease in oscillatory gamma power contrasts with previous studies in experienced meditators [1, 4, 8, 41, 51], which reported an increase in gamma both globally and locally in the frontocentral and posterior regions. A likely explanation lies in the fact that these studies did not account for possible modulations of non-oscillatory components of the signal. Indeed, the use of power spectral density as a measure of EEG or MEG oscillations requires caution [51], as the power spectrum reflects not only oscillatory (periodic) brain activity but also the level of non-oscillatory (aperiodic) brain activity, which is captured by the 1/f slope of the spectrum [15]. Of course, in the absence of significant changes in the aperiodic component, estimating oscillatory power with or without correcting for the 1/f slope, would not have much impact. However, as shown by our analyses, both meditation techniques showed a statistically significant reduction (i.e. flattening) of the slope compared to resting state. While these results do not falsify previous observations, they do suggest that the reported increase of gamma power in other studies may be largely due to an upward broadband shift across higher frequencies, rather than an increase in actual gamma oscillation amplitude. The disambiguation between the periodic and aperiodic changes induced by meditation in the high frequency range is crucial when it comes to the mechanistic interpretation. Indeed, the discrepancies we found between the corrected and uncorrected spectral power results point towards the relevance of considering the aperiodic component (i.e. the 1/f slope) of the power spectrum as a potentially important brain feature that is altered by meditation. At minimum, our results suggest that all future investigations of spectral power modulations under meditation should examine the effect of removing the aperiodic component.

To the best of our knowledge, there is only one meditation study to date that has investigated changes in the slope of the power spectrum [61]. The authors compared EEG spectral modulations associated with meditation and mind wandering between experienced meditators and a control group without meditation experience. Their findings indicate that experienced meditators showed a non-significant increase in the steepness of the 1/f slope (i.e. more negative values) during uninterrupted breath-focused meditation compared to rest. This discrepancy between their results and ours could be attributed to methodological parameters including their use of a 19-channel EEG cap and their focus on a more limited frequency range (2-30 Hz) for EEG power spectrum, as well as potential discrepancies in the parameters used for the estimation of the spectral slope.

A tentative hypothesis for the observed reduction in the 1/f slope across the frontal, parietal, occipital, and temporal regions is that it may reflect alterations in the excitation-inhibition (E-I) ratio. Indeed, modeling work suggests that the aperiodic slope reflects changes in E:I balance [20] with a reduction in the 1/f slope (i.e. its scaling exponent) reflecting an increase in the E:I ratio. Interestingly, we observed a stronger drop in the 1/f slope for Vipassana as compared to Samatha. Moreover, these changes in the scaling behavior of the power spectra are linked to changes in self-similarity, which in turn can be related to shifts in neural criticality.

### Meditation effects on complexity and criticality-related measures

The DFA analysis revealed prominent statistically significant decreases in the LRTC scaling exponent during Samatha and Vipassana meditations within gamma1 (30-59 Hz) and gamma2 (60-90 Hz) frequency bands. A reduction in LRTC generally suggests a decrease in the persistence or memory of the system (reduced information propagation over time), suggesting changes in the underlying dynamical processes or a shift towards more random or less correlated neural activity. Irrmischer et al. [30] showed that meditation practitioners had weaker LRTC during a focused attention meditation compared to rest, based on DFA analysis of EEG data. Interestingly, this reduction was not seen in participants with no meditation experience. The authors interpreted the results as a shift toward a subcritical regime and argued that the reduced autocorrelation within the signal may be associated with fewer distractions from the task and interpreted as a down-regulation of certain mental processing activities. Similarly, Walter and Hinterberg [76] found that experienced meditators showed a significant reduction in the broadband DFA exponent under three different meditation conditions (FAM, presence monitoring and thoughtless emptiness) compared to the resting state. Our results are not only consistent with the previous findings but also suggest that the changes in LRTC exponents for the high frequency envelopes is the most prominent difference between meditation and rest. Indeed, examining the feature importances obtained from a multi-feature Random Forest classifier identified the LRTC scaling exponent computed from the gamma1 and gamma2 signal envelopes (mainly in the regions belonging to the Frontoparietal, Default Mode, Dorsal and Ventral Attention networks) to contribute most to the successful classification. The reduction in LRTC scaling exponents during meditation suggests diminished temporal redundancy, which is posited to improve processing efficiency. [27]. This improved efficiency, which may lead to a more vivid conscious experience of the object of focus was observed when examining the gamma envelope, which plays a crucial role in attentive processing [32] and conscious perception [49]. Interestingly, we also found that the number of years of meditation expertise was positively correlated with the meditation-induced change in 1/f slope and gamma LRTC exponent.

To further explore changes in criticality, we analyzed neuronal avalanches, which at the critical point should exhibit scale invariance and obey the crackling noise relation. Only two studies have previously explored neuronal avalanches in the context of meditation, with differing results and interpretations [16, 76]. Neither of these studies examined the DCC, which is thought to be a more reliable indicator of criticality than solely power laws and their critical exponent values [44, 71]. Our finding of a decrease in DCC from both resting-state and Samatha to Vipassana in experienced meditators suggests that Vipassana meditation involves a shift of brain dynamics towards the critical point. No such shift was observed from rest to Samatha meditation. Interestingly, as an OM-type meditation style, Vipassana involves an open, unfocused attention to the body and of one’s surroundings, reminiscent of the high sensitivity to perturbations of the critical state. Meanwhile, Samatha meditation requires a focused attention on a specific object, such as the breath or a bodily sensation, which is consistent with a subcritical, input-dampening regime. Our findings thus highlight a dissociation between different meditation styles that aligns the phenomenology of these meditation states with the computational properties of the corresponding dynamical regimes.

The overall increase of the complexity index, Higuchi Fractal Dimension and Spectral Entropy in both Samatha and Vipassana is in line with recent literature. For meditation-naive subjects performing FA meditation, Lu and Rodriguez-Larios [40] report an increase in LZC, Sample Entropy (SE), and HFD compared to mind wandering. Although differences in HFD were widespread across electrodes, effects in LZC and SE were more pronounced in central electrodes. In experienced meditators, Walter and Hinterberg [76] found higher neuronal complexity during emptiness and focused attention as compared to resting with eyes closed, as captured by the SE and HFD values. Kakamanu et al. [34] analyzed EEG data of meditators with different levels of expertise in Vipassana practice and found increased HFD and permutation entropy in teachers and novices. An increased fractal dimension, as determined by Sevcik’s method, was also found in a calming meditation task by Vysata et al.[75]. Only the study by Young [80] found a decrease in LZC in highly experienced meditators performing different style of meditation practices as compared to mind wandering. The inconsistency of the results could be related to differences in meditation practices and their use of a 16 channels EEG system.

The increase in complexity during meditation observed here in source space MEG (and other studies using scalp EEG) may be related to a larger repertoire of subjective experiences during meditation. The entropic brain hypothesis [11, 10] explores the connection between entropy and consciousness, suggesting that a rich altered state of consciousness, such as those induced by psychedelics, may results from an enrichment effect of neuronal dynamics, reflected by increased entropy in brain data [3]. According to this theory, the degree of entropy in spontaneous brain activity captures the *richness* of conscious experience; for example, after consuming psychedelics, EEG signals and subjective experiences become more complex and disorganised [70]. Our findings indicate that the meditative states of Samatha and Vipassana, much like the altered states induced by psychedelics, generate greater neural complexity than is observed in the resting state. This similarity points to a potentially shared mechanism in how these different practices influence brain activity.

Our findings complement and extend those of D’Andrea et al. [17] who investigated the same dataset through the lens of criticality by using a microstate approach at ROI level and found that in both meditation conditions, the Hurst exponent values were reduced with respect to the RS condition, while the LZC values increased from rest to Samatha to Vipassana.

The present study has a number of limitations, which can hopefully be overcome in future experiments. First, our work is limited by the relatively small sample size, which is due to the rarity of participants with such extensive meditation expertise. Additionally, the potential confounding effect of age on the correlation between features and expertise warrants caution in interpretation. Furthermore, despite its widespread application, the Lempel-Ziv Complexity metric used here has a number of limitations which may be overcome by exploring alternative measures [50]. Lastly, the lack of a control group limits our ability to explore trait effects among experienced meditators with greater specificity.

## Conclusions

In summary, our research reveals that both forms of meditation studied lead to significant changes in brain dynamics. These include a leveling of the 1/f slope, a reduction in the high-frequency LRTC exponent, and increases in Lempel-Ziv Complexity and Spectral Entropy. These changes hint at an increased excitation-inhibition ratio during meditation. Contrary to previous findings, we observed a decrease in gamma power, which we believe is due to the correction of the power spectra by the 1/f slope. Moreover, the more years participants had practiced meditation, the greater the observed changes in the 1/f slope and gamma LRTC exponent during their meditation sessions. Using a Random Forest classifier, our analysis pinpointed the LRTC scaling exponent as a key factor in distinguishing between meditation and resting states. This investigation into the complexity and critical dynamics of the brain during meditation opens new paths for understanding the neuronal mechanisms at play in these states, suggesting that such measurements could be crucial for deciphering how meditation affects the brain.

## Acknowledgements

KJ was supported by funding from the Canada Research Chairs program and a Discovery Grant from the Natural Sciences and Engineering Research Council of Canada (NSERC). This work has been in part supported by a grant from the BIAL Foundation (Portugal) on the project “Mindfulness Meditation state and trait through the eyes of brain computational modeling,” grant number 80/20. AP was partially supported by the Short Term Mobolity grant of CNR and by the Gruppo Nazionale pe il Calcolo Scientifico (GNCS-INDAM).

## Authors contributions

A.P.–Software, Visualization, Methodology, Conceptualization, Writing – Original Draft. D.M.–Software. T. L.–Software. J.O’B.-Software, Methodology, Writing–Original Draft. P.T.-Software, Methodology. A.R.-Investigation. R.G.-Investigation. V.P.-Investigation. L.M.-Funding acquisition, Investigation. K.J.-Methodology, Conceptualization, Writing - original draft, Supervision, Project administration.

