## Supplementary for "Meditation induces shifts in neural oscillations, brain complexity and critical dynamics: Novel insights from MEG"

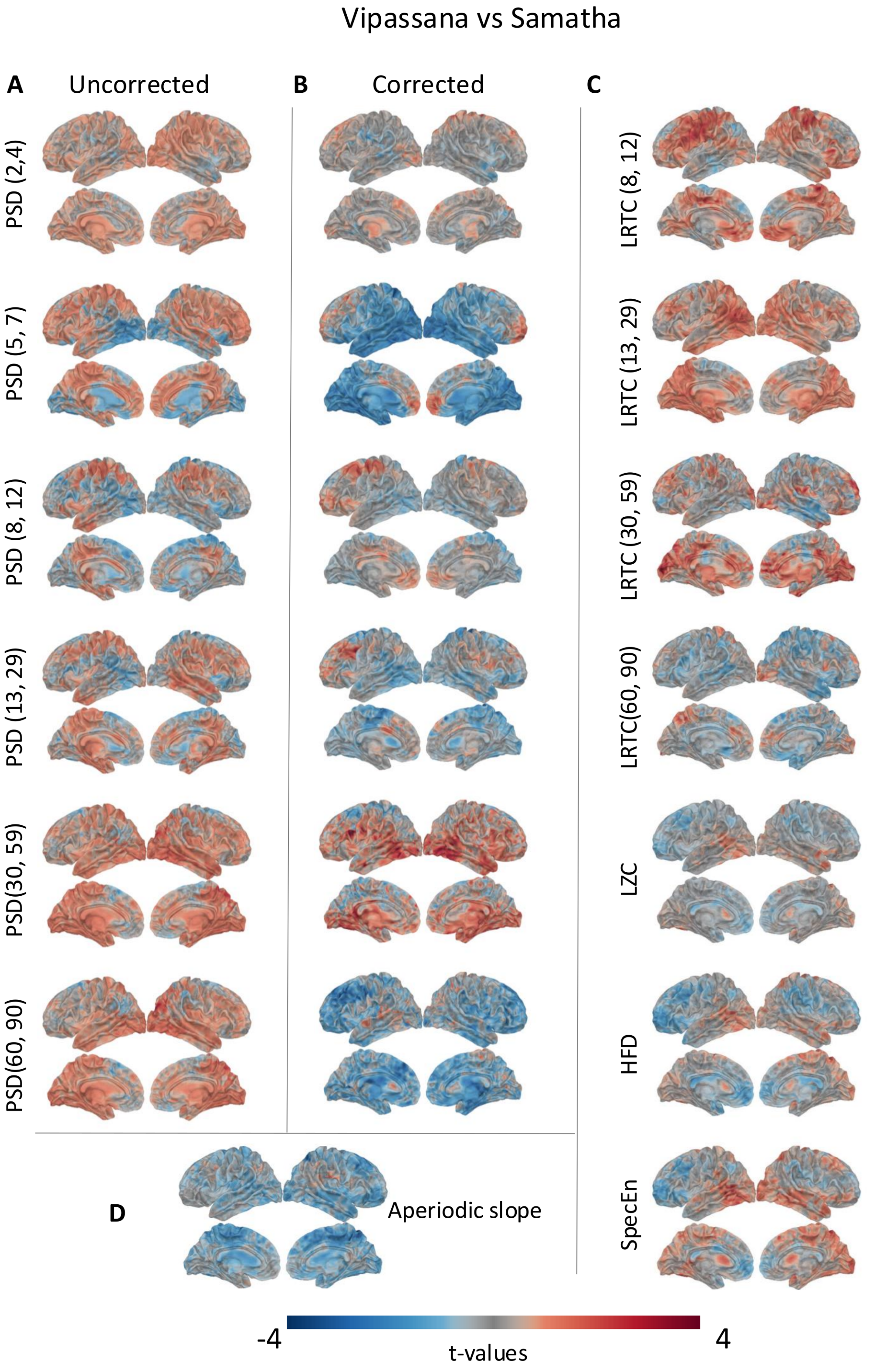


Figure S1: T-maps and results of cluster-based permutation test for all spectral, complexity and criticality related features for Vipassana as compared to Samatha. Although the t-maps show trends in the data, none of the effects survive corrections for multiple comparisons.

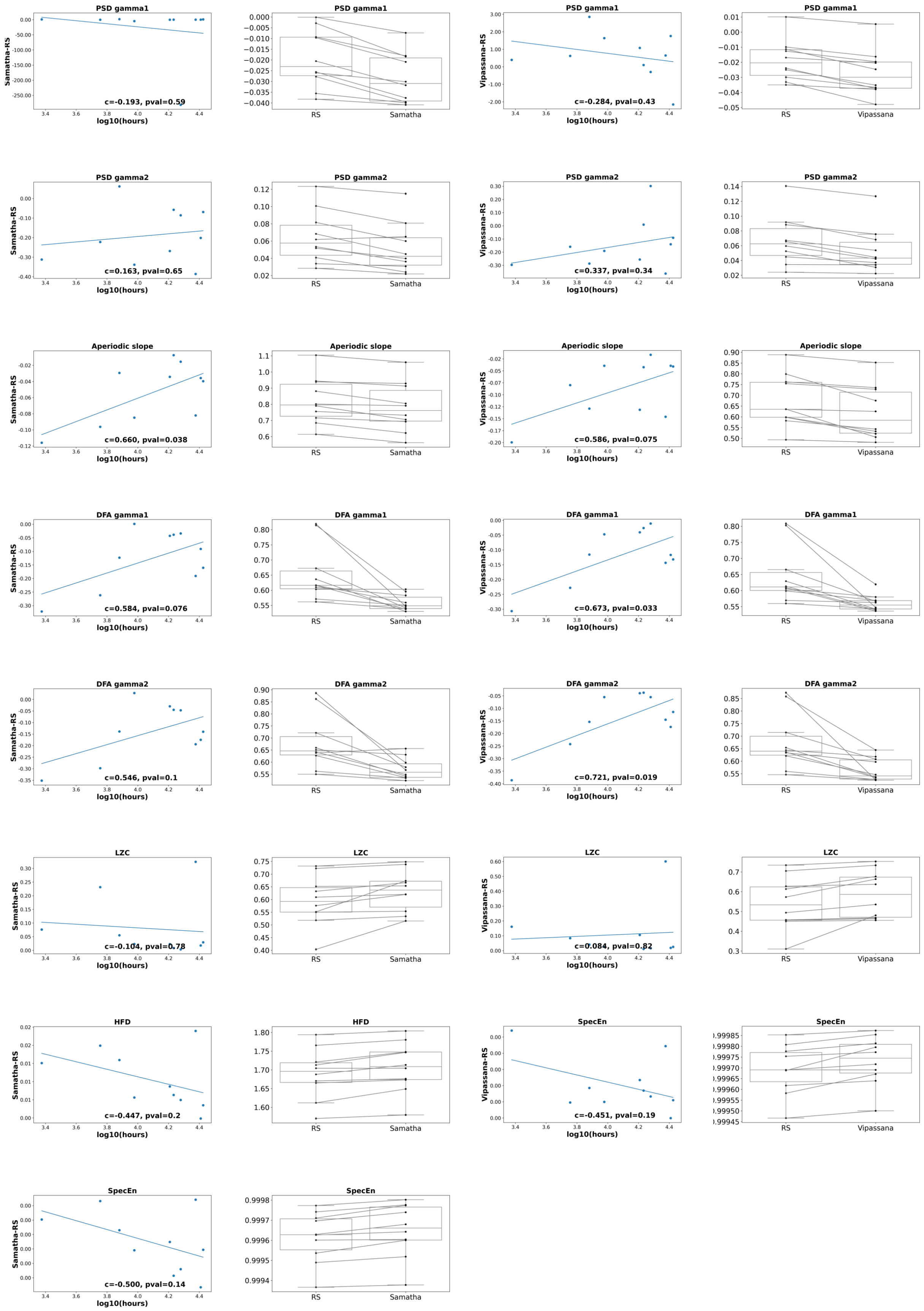


Figure S2: Correlations with hours of meditation and box plots, by feature The first and third columns show the Pearson correlation between hours of meditation and the features in which we found a state effect of Samatha and Vipassana, respectively. The second and fourth columns illustrate the box plot of individual features values (averaged within the significant clusters) for Samatha and Vipassana, respectively.

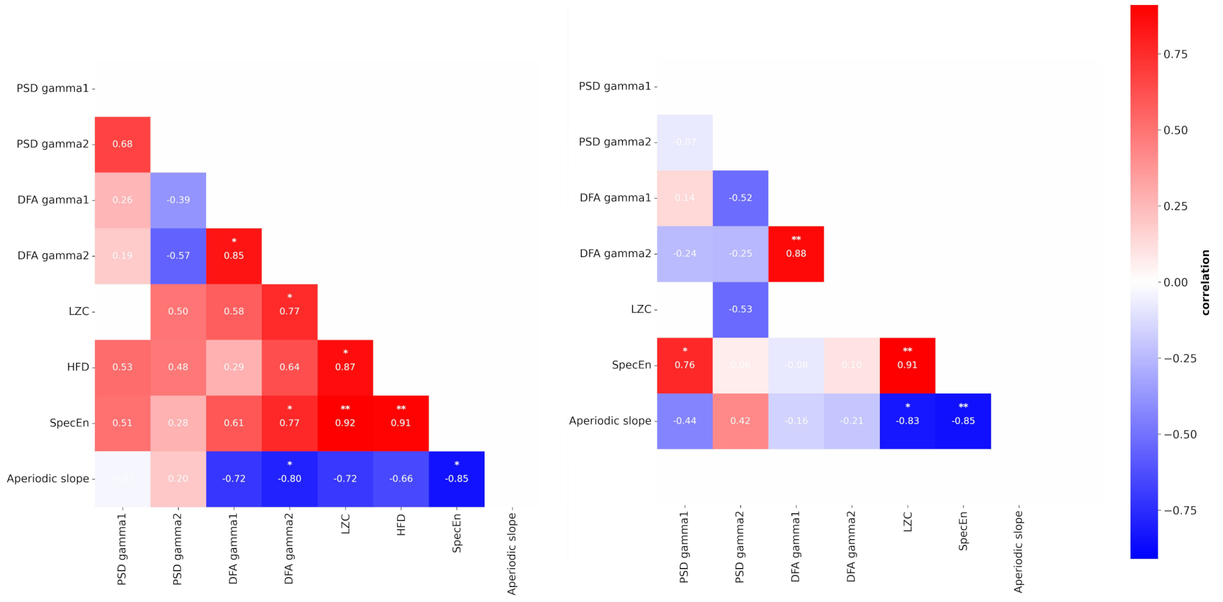


Figure S3: Correlations between the average values of each feature pair across common voxels in significant clusters

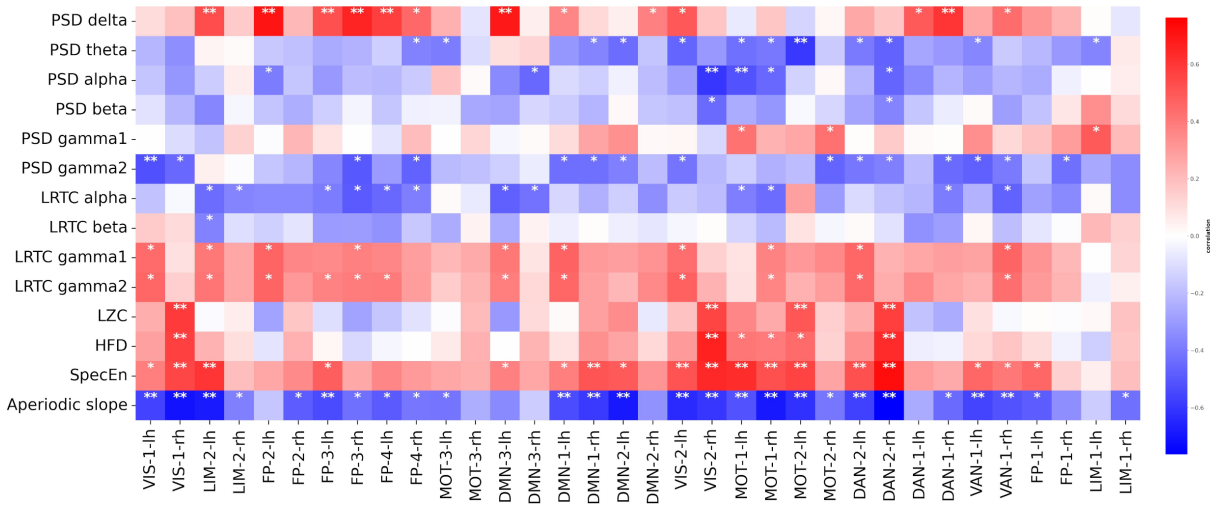


Figure S4: Correlations between DCC and all other features, computed across all conditions. * indicate $p < 0.05$
